# Epigenetic Patterns of *Xylanibacter ruminicola* in Bovine Rumen Across Seasons and Pregnancy

**DOI:** 10.64898/2026.06.07.730735

**Authors:** Ziming Chen, Chian Teng Ong, Marina R. S. Fortes, Kieren McCosker, Milou H. Dekkers, Alana C. Boulton, Basil S. Firewski, Elizabeth M. Ross

## Abstract

Ruminants obtain nutrients through the microbial fermentation of plant material in the rumen. *Xylanibacter ruminicola* is a highly abundant bacterial species in the rumen. During fermentation, *X. ruminicola* utilizes diverse carbohydrates from plant materials to synthesize propionate, a volatile fatty acid providing energy to ruminants. However, variation in pasture quality (e.g, nutrient and fibre content) across seasons and host pregnancy status can alter the rumen microenvironment, potentially affecting microbial activity. Bacteria in culture display distinct methylation (a reversible epigenetic modification capable of gene regulation) changes in response to the growth environment. We hypothesized that the changes to the rumen environment would affect the DNA methylation patterns within the *X. ruminicola* genome. Rumen fluid from 37 female Brahman cattle (17 pregnant) were sampled across four seasons. DNA methylation profiles (N6-methyladenine, N4-methylcytosine, and 5-methylcytosine) of *X. ruminicola* across seasons (varying pasture quality) and pregnancy statuses characterized using Oxford Nanopore sequencing. After correcting for relative abundance, DNA methylation levels within the coding DNA sequences of several *X. ruminicola* genes differed between seasons and pregnancy status. Most of these genes were classified as ExbD/TolR family proteins and related to the protein transport process. Our study demonstrates that the DNA methylation profiles of rumen *X. ruminicola* genes vary with host environment factors. These results provide insight into the role of bacterial DNA methylation in mediating interactions between bacteria and their environments.

**Lay Summary:** Ruminants rely on rumen microbes to convert plants into nutrients. *Xylanibacter ruminicola* is a bacterial species in the rumen that produces nutrients for ruminants during plant fermentation. Changes in pasture quality and host pregnancy status can influence the activity of *X. ruminicola*, as reflected in the DNA methylation profile across its genome. DNA methylation is a reversible DNA modification that can affect gene activity and help bacterial adaptation to changing environments. Rumen fluid from 37 female Brahman cattle (17 pregnant) across four seasons were used to evaluate the pasture quality and host pregnancy effects on the DNA methylation profile of *X. ruminicola*. The relative abundances of *X. ruminicola* were influenced by pasture quality, but not by host pregnancy. However, host pregnancy status and changes in pasture quality influenced the DNA methylation signatures of *X. ruminicola*. Genes with DNA methylation changes were associated with the protein transport process. These findings suggest that the DNA methylation profiles of rumen *X. ruminicola* vary with host environment factors.

**Teaser Text:** This study demonstrates that the DNA methylation profiles of rumen *Xylanibacter ruminicola* vary with host environment factors. These findings provide insight into the role of bacterial DNA methylation in mediating interactions between rumen bacteria and their environment.

## Introduction

*Xylanibacter ruminicola*, previously known as *Prevotella ruminicola* [1] or *Bacteroides ruminicola* [2], was first isolated from the bovine rumen in 1958 [2]. *Xylanibacter ruminicola* is a highly abundant bacterial species within the rumen microbial community [3, 4], playing a key role in nitrogen and carbohydrate metabolism [5-7]. Despite its inability to degrade cellulose, *X. ruminicola* can utilize various carbohydrates [7], including hemicelluloses [8] and cellobiose [9]. In symbiosis with other rumen microbes, *X. ruminicola* can hydrolyze plant materials into monomers, followed by the synthesis of volatile fatty acids (VFAs), CO_2_, and H_2_ [10]. The VFA metabolites generated during microbial fermentation, such as propionate and butyrate, supply nearly 80% of energy for ruminants [11]. In addition, propionate synthesized by *X. ruminicola* can serve as a hydrogen sink, reducing methane emissions [12]. In return, ruminants provide an ideal habitat, the rumen, for microbial growth, constituting microbe-host interactions in the rumen ecosystem.

Although the rumen is a relatively stable ecosystem, the rumen microenvironment is highly dynamic and can potentially be influenced by external and internal factors, such as feed quality and host physiological status. For instance, the quality of forage grass varies with different environmental factors across seasons, leading to changes in nutrient and fibre contents [13, 14] available to *X. ruminicola* and other rumen microbes. In addition, the host pregnancy status can influence gut microbial structures and metabolism to enhance energy storage efficiency, supporting fetal development [15, 16]. To maintain the stability and performance of the rumen ecosystem, *X. ruminicola* and other microorganisms must rapidly adapt to these dynamic and complex changes.

DNA methylation is a reversible modification of DNA molecules. In batch culture, DNA methylation patterns shift in response to environmental changes [17] presumably as an adaptive response. Bacteria exhibit diverse DNA methylation forms: 5-methylcytosine (m5C), N4-methylcytosine (m4C), and N6-methyladenine (m6A) [18]. In bacteria, DNA methylation not only protects the bacterial genome from viral invasion but also regulates gene expression [18, 19]. Reversible gene regulation mediated by DNA methylation may enable bacteria to respond rapidly to transient environmental changes [20, 21], suggesting that DNA methylation profiles may vary during microbial adaptation. However, variation in microbial DNA methylation profiles across environmental conditions remains largely unknown, partly due to the limited resolution of conventional DNA methylation detection methods. This limitation is overcome with new sequencing techniques (e.g. Oxford Nanopore Technologies; ONT) that can simultaneously identify various forms of DNA methylation to characterizee diverse DNA methylation patterns in prokaryotes, such as DNA methylation motif identification [22] and characterizing DNA methylation variation at the single-nucleotide level [23].

In this study, we hypothesized that methylation patterns of *X. ruminicola* would change in response to rumen environmental conditions induced by dietary changes across seasons and physiological changes in pregnancy. To test this hypothesis, DNA methylation profiles of *X. ruminicola* in the bovine rumen were extracted from microbiome data from 37 female Brahman cattle across seasons and pregnancy status.

## Materials and methods

### Animal ethics and sampling

Animals used in this study were Brahman cattle (*Bos taurus indicus*) obtained from a commercial supplier in Queensland, Australia, and accommodated at the Queensland Animal Science Precinct (QASP) - the University of Queensland Gatton campus. Animals were fed pasture hay, which was collected and delivered to Feed Central in Queensland, Australia, for feed quality assessment. The collection of bovine rumen samples was conducted under the Animal Ethics Number 2022/AE000438, the University of Queensland Animal Ethics Committee. The sample collection in this study was performed from winter 2023 to autumn 2024 in Queensland, Australia.

For rumen fluid collection, the stomach tube was guided into the rumen through the secure mouth gag. Upon reaching the rumen, the stomach tube was attached to the top of a polypropylene vacuum flask connected with a manual vacuum pump. Around 50 ml rumen fluid was initially pumped into the vacuum flask and discarded to minimize the saliva contamination. The other 50 ml rumen fluid was pumped and transferred into a sterile 50 ml Falcon tube (Eppendorf, Germany). Rumen fluid was further aliquoted into 2 ml tubes with 1.5 ml fluid per tube. The tube was temporarily maintained on ice and subsequently stored at −80°C.

### DNA extraction and sequencing

The rumen fluid sample (1.5 ml) was centrifuged at 14,000 rpm for 5 minutes at 4°C, followed by supernatant removal. DNA was extracted from the remaining pellets using the QIAamp PowerFecal Pro DNA Kit (QIAGEN, Germany) following the manufacturer’s instructions. The ONT Native Barcoding Kit 96 V14 (ONT, UK) was used for DNA library preparation according to the manufacturer’s protocol. The DNA library was loaded on a FLO-PRO114M Flow Cell (R10.4.1), followed by sequencing on a PromethION 2 Solo (ONT, UK).

### Quality control, alignment, and taxonomic classification

DNA data were basecalled by ONT Dorado v1.1.1 using the super-accurate model (SUP), with DNA methylation detection function enabled (5-methylcytosine (m5C), N4-methylcytosine (m4C), and N6-methyladenine (m6A)). To remove the host data, raw reads were mapped to the bovine reference genome (GCF_002263795.3) using minimap2 v2.28 [24], followed by the extraction of unmapped reads, denoted as de-host reads, using SAMtools v1.18 [25]. De-host reads shorter than 250 bp and average Phred quality score lower than 10 were removed by chopper v0.10.0 [26].

Filtered reads were aligned to the *X. ruminicola* reference genome (GCF_000025925.1) using minimap2 v2.28 [24] with the flag “-y” to preserve the DNA methylation information. Primarily mapped reads were extracted using SAMtools v1.18 [25]. To minimize the effects of genetic similarity among species on taxonomic classification, a standard protein database (as of 11 Feb 2026) with complete RefSeq for bacteria, archaea, viruses, humans, and known plasmids was constructed using Kraken2 v 2.17.1 [27]. Primary mapped reads were taxonomically classified against the standard protein database using Kraken2 v2.17.1 [27], followed by the extraction of reads classified to *X. ruminicola* using KrakenTools v1.2.1 [28]. The numbers of reads and bases for filtered and extracted data were calculated using a custom bash script. To minimize the read length effects, the relative abundance of *X. ruminicola* was defined as the division of extracted sequence bases by filtered bases.

### Effects of seasons, grass quality, and pregnancy on relative abundance

The effects of seasons and pregnancy statuses on the relative abundances of *X. ruminicola* were evaluated using a linear mixed model as follows:

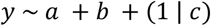

where *y* is the relative abundance of *X. ruminicola*; *a* is the pregnancy information of an animal; *b* is the sample collection season; *c* is the animal identifier. The linear mixed model was weighted by the total base number and fitted using bound optimization by quadratic approximation (BOBYQA). The constructed linear mixed model was evaluated using analysis of variance (ANOVA). The *P-value* threshold was 0.05.

To evaluate the effects of grass starch and protein contents on the relative abundances of *X. ruminicola*, a linear mixed model was constructed as follows:

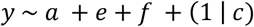

where *y* is the relative abundance of *X. ruminicola*; *a* is the pregnancy information of an animal; *e* is the crude protein contents; *f* is the starch contents; *c* is the animal identifier. The linear mixed model was weighted by the total base number and fitted using BOBYQA. The constructed linear mixed model was evaluated using ANOVA. The *P-value* threshold was 0.05.

### DNA methylation calling on each genomic locus

The BAM file and the reference genome were processed with ONT modkit v0.4.5 to generate a bedMethyl file with DNA methylation information at each genomic position. The global modification calling threshold was set to 0.9 (--filter-threshold 0.9) to minimize the effects of low sequencing coverage and increase the DNA methylation calling reliability. For each position in the bedMethyl table, the locus methylation level was defined as the number of methylated bases (N_mod_) divided by the total number of bases (N_valid_cov_). The locus DNA methylation proportions were used for principal component analysis (PCA).

### De novo DNA methylation motif detection

*De novo* motif identification was performed using ONT modkit v0.4.5 and Snappy [29]. For ONT modkit v0.4.5, rare DNA methylation motifs in each sample were identified using the reference file and corresponding locus-based bedMethyl table, following the settings in our previous study with slight modification [30]. In brief, the minimum coverage was set to 5x, and the minimum site required for a motif was 25. For Snappy [29], the bedMethyl file from ONT Modkit v0.4.5 and the reference genome were used for identifying motifs.

### DNA methylation analysis on gene bodies

The coding DNA sequences (CDSs), denoted as gene bodies, were retrieved from the *X. ruminicola* RefSeq GFF annotation file (GCF_000025925.1) and converted into a tsv file using AGAT v1.4.1 [31]. The tsv file was filtered with the “source_tag” as “Refseq” and “primary_tag” as “gene” using a bash script. Genomic positions within the gene bodies were used for DNA methylation analysis. The effects of seasons and pregnancy statuses on the DNA methylation level of a specific genomic position were evaluated using a generalized linear mixed-effects model as follows:

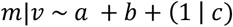

where *m* is the DNA methylation level of a specific genomic position; *v* is the valid sequencing depth at a specific genomic position; *a* is the pregnancy information of an animal; *b* is the sample collection season; and *c* is the animal identifier. *m*|*v* is the binomial response of DNA methylation levels, weighted by variation in sequencing depth among samples. The generalized linear mixed-effects model was fitted using BOBYQA. The constructed linear mixed model was evaluated using ANOVA. The *P-value* was adjusted by Bonferroni correction, grouped by fixed effects and DNA methylation forms. The adjusted *P-value* threshold was 0.05.

To evaluate the effects of grass starch and protein contents on the DNA methylation profile in *X. ruminicola*, a generalized linear mixed model was constructed as follows:

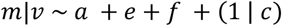

where *m* is the DNA methylation level of a specific genomic position; *v* is the valid sequencing depth at a specific genomic position; *a* is the pregnancy information of an animal; *e* is the crude protein contents; *f* is the starch contents; *c* is the animal identifier. *m*|*v* is the binomial response of DNA methylation levels, weighted by variation in sequencing depth among samples. The generalized linear mixed-effects model was fitted using BOBYQA. The constructed linear mixed model was evaluated using ANOVA. The *P-value* was adjusted by Bonferroni correction, grouped by fixed effects and DNA methylation forms. The adjusted *P-value* threshold was 0.05.

### Functional annotation analysis

Genomic positions with DNA methylation levels affected by seasonal, grass quality or pregnancy changes were extracted, followed by the ID retrieval of corresponding genes. The NCBI Database for Annotation, Visualization, and Integrated Discovery (DAVID) were used for functional annotation analysis by uploading the extracted Gene IDs. The *P-value* threshold after False Discovery Rate (FDR) correction was 0.05.

## Results

Rumen fluid microbial samples were taken from 37 female Brahman cattle (17 of which were pregnant) fed pasture hay and accommodated in the Queensland Animal Science Precinct (QASP) - the University of Queensland Gatton campus. A full description of the animal trial is detailed in [32]. Samples were collected across four seasons within a year for each animal (from winter 2023 to autumn 2024), followed by DNA extraction and Oxford Nanopore sequencing. A total of 148 samples were collected from four seasons for Nanopore sequencing, generating 351 Gb sequence data after quality filtering. Of the filtered data, 3.83 ± 1.52% of bases aligned to the *X. ruminicola* reference genome and were used for taxonomic classification. These filtered reads were used to evaluate variation in relative abundance and DNA methylation across seasons and pregnancy statuses.

### Relative abundances of X. ruminicola varied across seasons

Species within the same Order level can share high genetic similarity [33], causing the misalignment of sequences from different species and the misrepresentation of relative abundance. However, small sequence changes can shift open reading frames; thereby, using a translated search enables higher sensitivity in taxonomic classification. Accordingly, reads primarily mapped to the *X. ruminicola* genome (14 Gb total, 1.8M reads) were translated and searched against the Kraken2 [27] protein database, followed by the extraction of reads classified to *X. ruminicola*. This step removed 85.20% of reads likely incorrectly aligned to the *X. ruminicola* genome.

Due to variation in read length across samples (Figure 1A), the relative abundance of *X. ruminicola* was calculated using the number of bases instead of read numbers. A linear mixed model was used to evaluate the seasonal and pregnancy effects on the relative abundance of *X. ruminicola*. The seasonal changes showed a significant effect on the relative abundance of *X. ruminicola* in the rumen (*P* < 0.05; Figure 1B). However, pregnancy status showed no significant effect on the abundance of *X. ruminicola* (*P* > 0.05).

**Figure 1.**
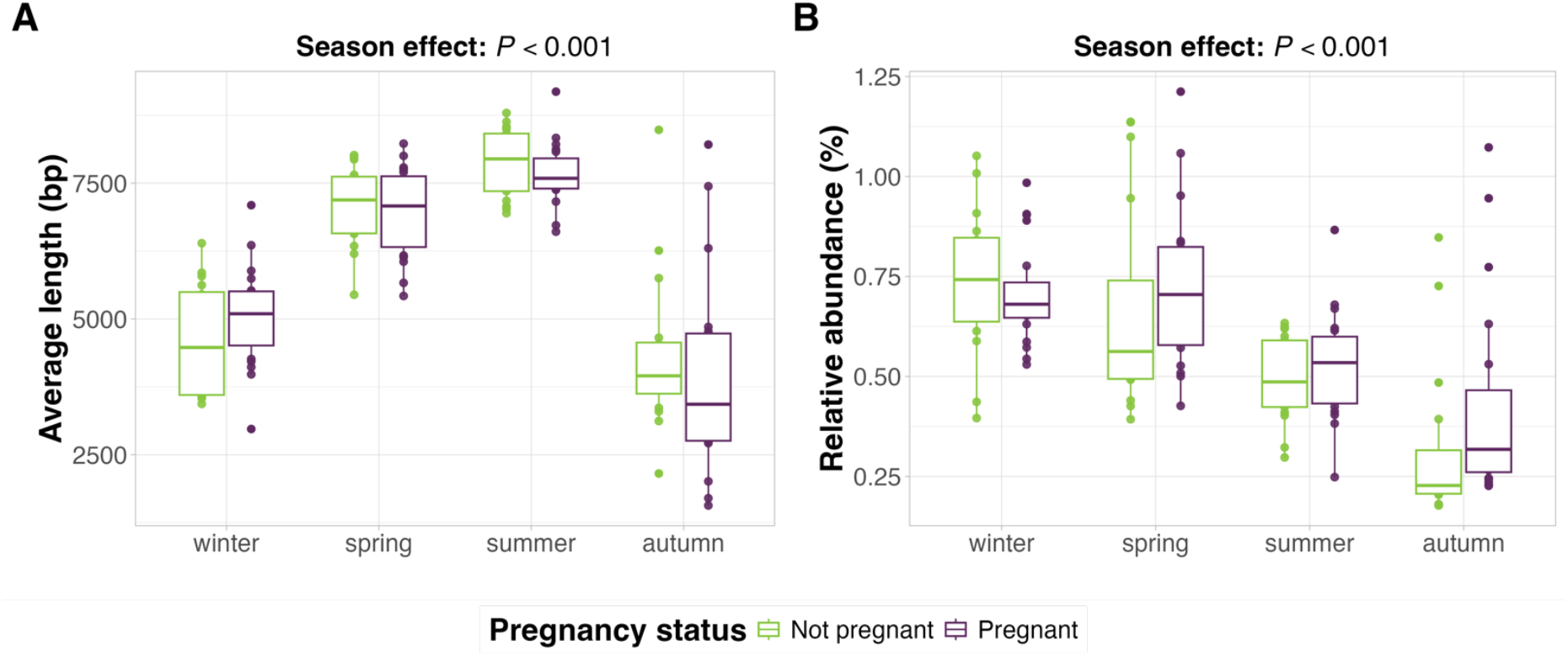
The average read lengths (A) and relative abundances (B) of *Xylanibacter ruminicola* associated reads across season and pregnancy. The effects of season and pregnancy on *X. ruminicola* relative abundance were evaluated using a linear mixed model and analysis of variance.

### Global DNA methylation patterns

DNA methylation is a gene regulatory element in organisms [34, 35]. We hypothesized that pregnancy status and seasonal changes may affect microbial activity, as indicated by shifts in DNA methylation signatures, such as DNA methylation motifs. Therefore, *de novo* motif identifications were first performed by Modkit v0.4.5 and Snappy [29] to examine variation in DNA methylation motif patterns across samples. However, no DNA methylation motifs were identified in any samples, possibly due to the relatively low sequencing depths.

A previous study [30] showed that locus-based DNA methylation calling can capture most DNA methylation variance. Therefore, we utilized the matrix of methylation proportions at each detected genomic position to visualize global DNA methylation variation across samples. However, no distinct seasonal or pregnancy-related clustering patterns were observed (Figure 2A). Similarly, using DNA methylation levels within coding DNA sequences (CDSs) showed no clustering pattern (Figure 2B).

**Figure 2.**
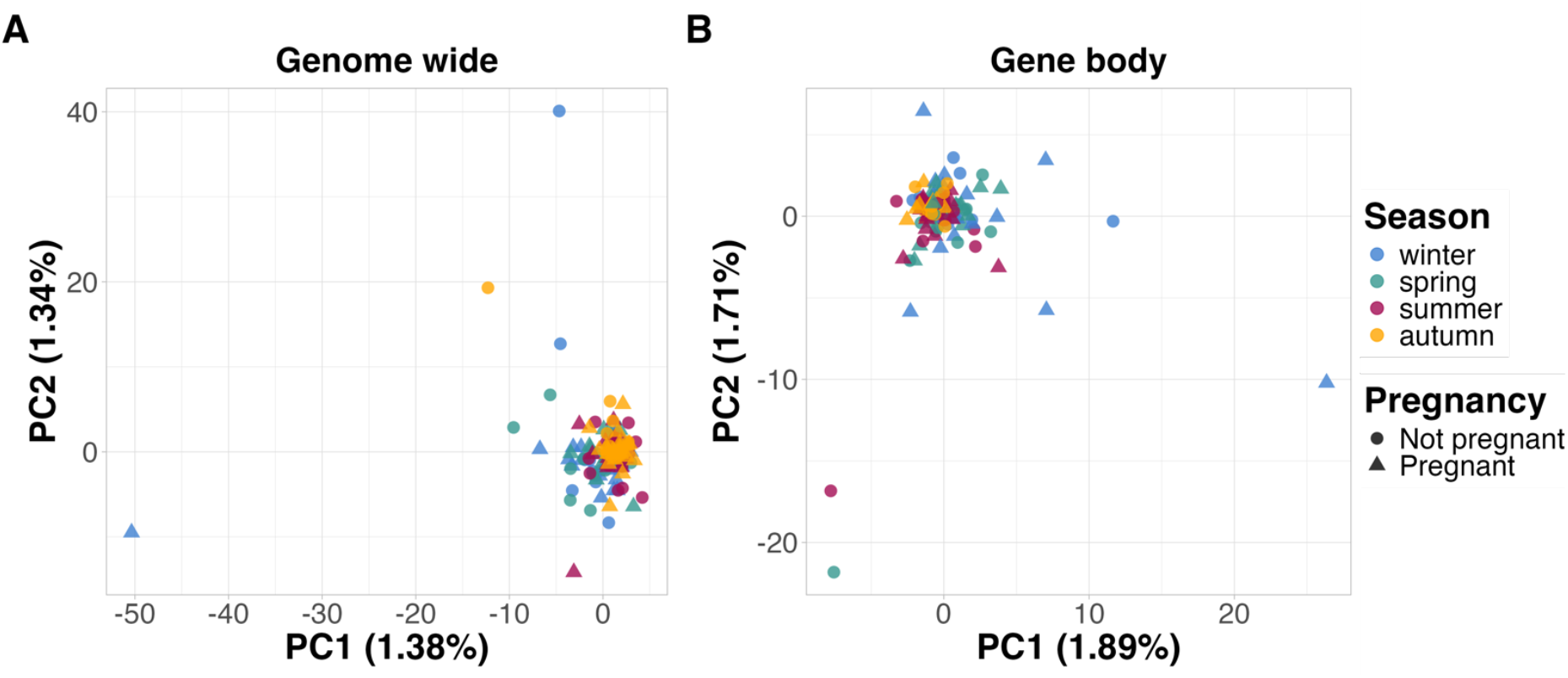
Principal component analysis using loci across the genome (A) and within the gene body (B) of *Xylanibacter ruminicola*. A matrix containing the DNA methylation level at each genomic locus with any methylation detected was used for principal component analysis. Methylation level was calculated as the proportion of methylated bases at the given locus in the given sample.

### DNA methylation levels of genes were affected by seasons and pregnancy

The rumen is a relatively stable microenvironment due to host homeostasis; therefore, the pregnancy status and seasonal changes may not significantly alter the global DNA methylation pattern of *X. ruminicola*. However, these environmental changes may affect the expression of several genes, reflected in DNA methylation variation in corresponding coding DNA sequences (also known as gene bodies). Therefore, the variation in DNA methylation levels of genomic loci within CDSs across seasons and pregnancy statuses was examined using a generalized linear mixed-effects model.

A total of 3,032 genes were tested in this analysis. After a Bonferroni correction, four genomic positions within gene bodies (corresponding to three distinct genes) exhibited seasonal variation in DNA methylation levels (Corrected-*P* < 0.05; Figure 3), two of which were also affected by pregnancy status. All genomic loci within gene bodies influenced by seasons or pregnancy exhibited variation in m4C levels (Table 1). These findings demonstrated that seasonal and pregnancy changes can cause DNA methylation shifts within the CDSs of genes in *X. ruminicola*.

**Figure 3.**
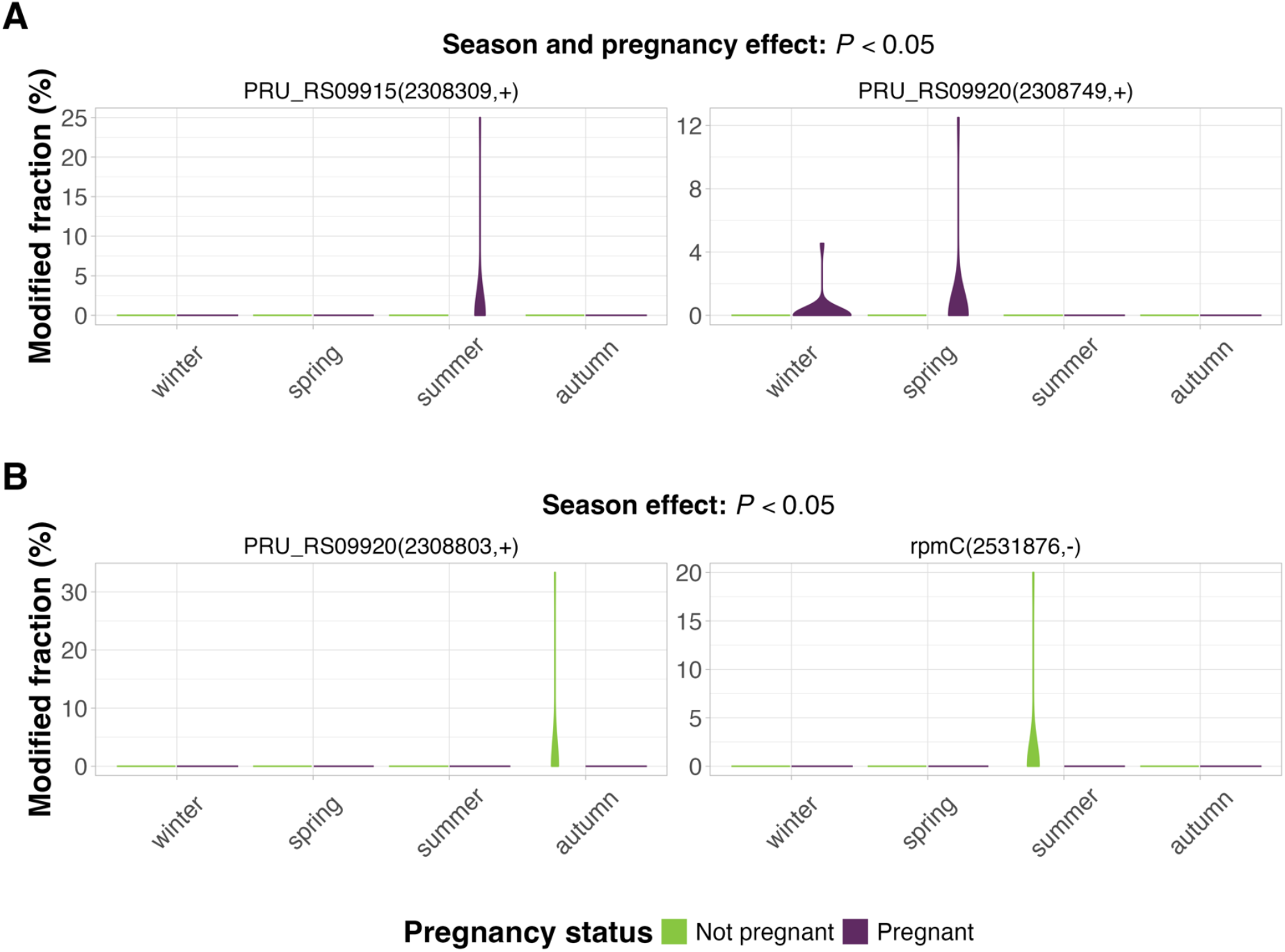
Genomic loci on chromosome NC_014033.1 with significant DNA methylation level changes. **(A) Genes with DNA methylation levels affected by both seasons and pregnancy. (B) Genes with DNA methylation levels affected by only seasons**. The strip label displayed above the plot indicates: gene symbol (genomic position, strand). The season and pregnancy effects on DNA methylation levels were evaluated through a generalized linear mixed model.

**Table 1.**
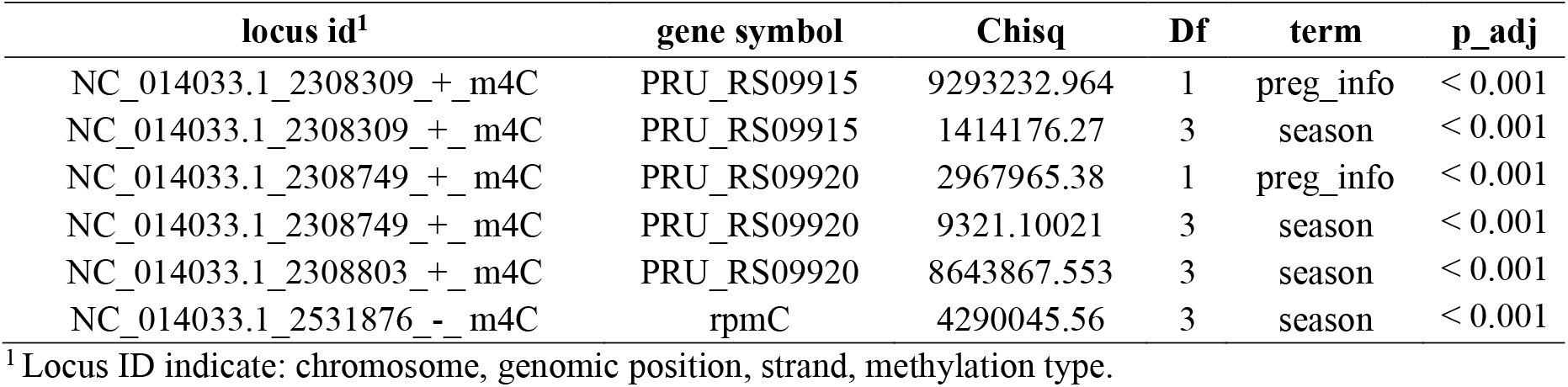
Evaluation of season and pregnancy effects on DNA methylation levels within gene bodies using ANOVA.

| locus id<sup>1</sup> | gene symbol | Chisq | Df | term | p_adj |
| --- | --- | --- | --- | --- | --- |
| NC_014033.1_2308309_+_m4C | PRU_RS09915 | 9293232.964 | 1 | preg_info | < 0.001 |
| NC_014033.1_2308309_+_m4C | PRU_RS09915 | 1414176.27 | 3 | season | < 0.001 |
| NC_014033.1_2308749_+_m4C | PRU_RS09920 | 2967965.38 | 1 | preg_info | < 0.001 |
| NC_014033.1_2308749_+_m4C | PRU_RS09920 | 9321.10021 | 3 | season | < 0.001 |
| NC_014033.1_2308803_+_m4C | PRU_RS09920 | 8643867.553 | 3 | season | < 0.001 |
| NC_014033.1_2531876_-_m4C | rpmC | 4290045.56 | 3 | season | < 0.001 |

<sup>1</sup> Locus ID indicate: chromosome, genomic position, strand, methylation type.

### Seasonal grass qualities affected X. ruminicola abundances and epigenetic patterns

Although the rumen temperature is relatively stable across seasons due to host thermoregulation, pasture hay quality varies across the year (Table 2). Therefore, we hypothesized that seasonal changes in the nutritional compositions of pasture can affect *X. ruminicola* microbial activity, reflected in the variation in relative abundances and DNA methylation. A generalized linear mixed-effects model was used to evaluate how pasture starch and protein contents influence relative abundances and DNA methylation of *X. ruminicola* with the pregnancy effect.

**Table 2.** Pasture hay quality across seasons.

| Month | Season | Avg. Daily Temp (°C) | Avg. Daily Rainfall (mm) | ME (MJ/kg)<sup>1</sup> | Crude Protein (%) | Starch (%) | NDF%<sup>2</sup> | ADF%<sup>3</sup> |
| --- | --- | --- | --- | --- | --- | --- | --- | --- |
| August | winter | 7.6 to 25.1 | 0.19 | 7.04 | 5.00 | 1.70 | 79.70 | 44.20 |
| November | spring | 16.7 to 29.3 | 5.9 | 6.95 | 5.30 | 1.00 | 79.80 | 47.70 |
| January | summer | 20.9 to 31.6 | 6.36 | 8.50 | 10.70 | 1.50 | 66.48 | 35.04 |
| March | autumn | 14.2 to 33.6 | 0.81 | 8.60 | 7.80 | 1.30 | 67.10 | 38.40 |

<sup>1</sup> ME: Metabolizable energy
<sup>2</sup> NDF: Neutral detergent fibre
<sup>3</sup> ADF: Acid detergent fibre

The relative abundance of *X. ruminicola* was significantly affected by the crude protein contents of pasture hay (*P* < 0.05; Table S3), but not starch composition. We then included genomic loci of 3,032 genes to evaluate the effects of starch and protein contents on the DNA methylation profiles of *X. ruminicola*. After a Bonferroni correction, nine genomic loci (corresponding to four distinct genes) showed differential methylation associated with pasture quality and pregnancy condition (Corrected-*P* < 0.05; Figure 4). Most differentially methylated genomic loci were m4C, followed by m6A (Table 3). Overall, seasonal variation in pasture hay quality can affect the relative abundances and DNA methylation profiles of *X. ruminicola*.

**Table 3.**
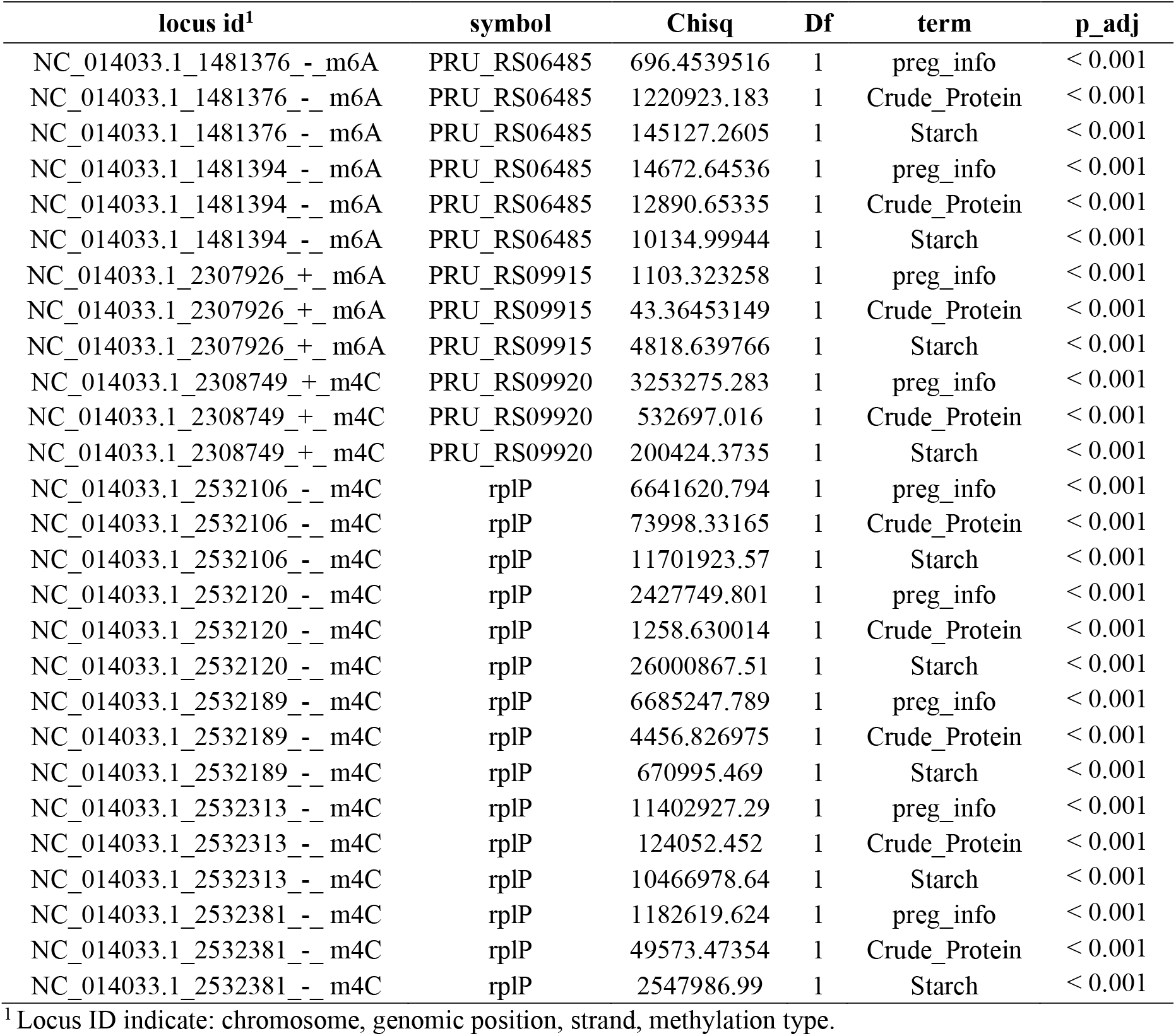
Evaluation of grass quality and pregnancy effects on DNA methylation levels within gene bodies using ANOVA.

**Figure 4.**
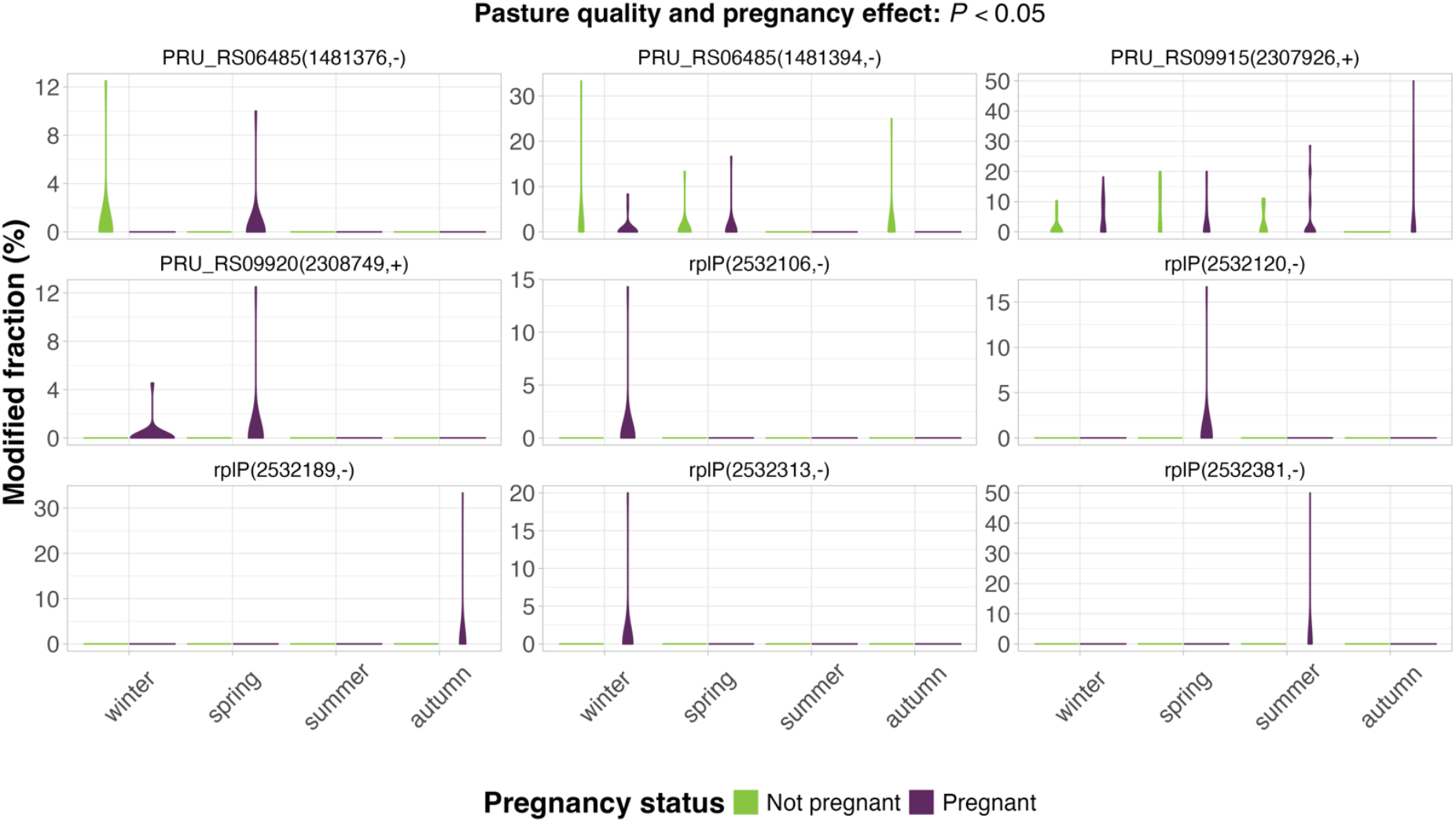
Genomic loci on chromosome NC_014033.1 with significant DNA methylation level changes between pasture quality and pregnancy. The strip label displayed above the plot indicates: gene symbol (genomic position, strand). The season and pregnancy effects on DNA methylation levels were evaluated through a generalized linear mixed model.

### Differentially methylated genes were related to protein transport

The functions of genes with DNA methylation shifts in relation to season, pasture quality, or pregnancy were examined. Genes (e.g., *PRU_RS09915* and *PRU_RS09920*) influenced by season, pasture quality, or host pregnancy were involved in protein transport biological process (*P* < 0.05; Table S4). These findings suggested that the seasonal quality changes in pasture and host pregnancy may affect the microbial activity of *X. ruminicola*, reflected in changes in DNA methylation levels of the corresponding genes.

## Discussion

This study characterized the relative abundance and DNA methylation patterns of *X. ruminicola* in the bovine rumen across seasons and pregnancy. Seasonal grass quality changes altered the abundance and DNA methylation profile of *X. ruminicola*, while host pregnancy only influenced its DNA methylation signature. Genes that showed differential methylation under varied grass quality and host pregnancy status were associated with protein transport. Our findings indicated that environmental changes may affect the microbial activity of *X. ruminicola*, reflected in variation in its relative abundance and DNA methylation profiles.

The community of *X. ruminicola* was influenced by seasonal variation in the quality of pasture. Environmental changes (e.g., temperature and precipitation [13, 14]) can alter the nutrient and fibre contents of various grass types [36] available to the rumen microbial community, which was consistent with our feed assessment of pasture hay. Under different nutrient compositions, microbes can exhibit different growth rates due to their metabolic capabilities and nutritional preferences [37, 38]. These differential growth rates can lead to shifts in microbial abundances within the rumen community [39]. Aligning with previous studies investigating the effects of seasonal quality variation of forage grass on rumen microbial community [40, 41], the relative abundance of *X. ruminicola* in the bovine rumen in our study also varied with pasture quality. These findings indicated that pasture or grass quality is one of the underlying factors affecting rumen microbial structures.

Seasonal changes in the nutrient composition of pasture hay may influence the nutrient uptake activity of *X. ruminicola*. Under nutrient variation, microbes must undergo proper gene regulation to support the stability and performance of the rumen ecosystem [42, 43]. A previous study found that the ExbD/TolR family protein in *X. ruminicola* showed differential expression in response to various polysaccharides and pectin [44]. Similarly, our study found that ExbD/TolR family protein genes in *X. ruminicola*, such as *PRU_RS09920* and *PRU_RS09915*, exhibited DNA methylation variation under varying pasture hay quality. The ExbD protein interacts with ExbB and TonB to form the TonB-ExbB-ExbD complex, also known as the Ton complex, which facilitates active nutrient transport [45]. In the Ton complex, TonB connects the inner membrane to the TonB-dependent transporters (TBDTs) on the outer membrane. Energy from the proton motive force of the cytoplasmic membrane is captured by ExbB and ExbD and transferred to TBDTs through TonB, enabling the active nutrient uptake into the periplasm [46]. Together, these findings suggest that the pasture hay can affect nutrient uptake activity in *X. ruminicola*, as reflected in the DNA methylation variation in genes related to the ExbD protein.

Host pregnancy may affect the microbial activity of *X. ruminicola* to enhance energy metabolism. A previous study reported enriched rumen microbial glucose biosynthesis and carbohydrate metabolism in pregnant sheep [47], indicating improved energy utilization during gestation. Our study found that DNA methylation levels of ExbD/TolR family protein genes were affected by pregnancy conditions, suggesting the potential regulation of their gene activity. As discussed above, ExbD is a crucial component of the Ton complex that facilitates active nutrient transport [45]. Therefore, the regulation of ExbD protein expression during gestation may enable the efficient nutrient utilization to alleviate the negative nutrient balance of maternal parents and support fetal development. However, whether the host pregnancy status affects the expression of ExbD protein requires further investigation through meta-transcriptomics or meta-proteomics.

Similar to previous studies using epigenetic signals in mammalian tissues [48], cell-type confounding influences microbiome DNA methylation analyses. In microbiome DNA methylation studies, this cell-type confounding is primarily caused by the high sequence similarity across microbial species, particularly for taxa within the same Order [33]. This cross-species sequence similarity can result in sequences from different species aligning to the same genome. Consequently, such misclassification interrupts the DNA methylation signals at a genomic locus, leading to erroneous biological inference. In our study, to minimize cell-type confounding effects, data were filtered using both alignment and taxonomic classification. Although a relatively abundant bacterial species, *X. ruminicola*, was selected, a high proportion of reads were excluded during data filtering, resulting in limited sequencing depth for subsequent DNA methylation analyses. This limited sequencing depth constrained our ability to accurately characterize the methylation level at each genomic locus. This suggests that sequencing depth remains a major issue even for relatively high-abundance microbial species. Therefore, we recommend that future studies first estimate the relative abundance of target species in the specific microbiome sample type before calculating the amount of data needed to generate sufficient sequencing depth of the target.

Our findings demonstrate that an analysis pipeline independent of the DNA methylation motif can provide extra biological information for microbiome samples, especially under relatively low sequencing coverages. Previous prokaryotic DNA methylation studies focused on *de novo* DNA methylation motif identification [49-53], which enables the evaluation of microbial evolution resulting from the frequent genetic transfer in nature [54-56]. However, due to the limitation of sequencing coverage, no specific DNA methylation motif was identified in this study. This indicates that higher sequencing depths are recommended for *de novo* DNA methylation motif identification for microbiome samples, especially for microbes with low relative abundances. However, using an analysis pipeline independent of DNA methylation motifs, we identified that DNA methylation levels within the gene bodies of several genes were affected by seasons, which is not achievable when relying on DNA methylation motif discovery. Overall, our findings demonstrate that investigating prokaryotic methylation patterns can provide insight into the molecular mechanisms that allow host-associated prokaryotes to rapidly adapt to changing environments.

## List of Abbreviations

ANOVA: Analysis of variance
BOBYQA: Bound optimization by quadratic approximation
DAVID: Database for Annotation, Visualization, and Integrated Discovery
FDR: False Discovery Rate
m4C: N4-methylcytosine
m5C: 5-methylcytosine
m6A: N6-methyladenine
ONT: Oxford Nanopore Technologies
PCA: Principal component analysis
SUP: Super-accurate model
TBDTs: TonB-dependent transporters
VFAs: Volatile fatty acids

## Availability of data and materials

Coding scripts were available on the GitHub repository: https://github.com/SimonChen1997/DNA_methylation_X.ruminicole_rumen_pregnancy_season

## Acknowledgements

We appreciate the support from Mr Tony Cavallaro during sample collection. We would like to thank the colleagues at the Centre for Animal Science (Queensland Alliance for Agriculture and Food Innovation, University of Queensland) for providing valuable feedback for this study. We acknowledge the support from the Australian Government Research Training Program Scholarship. We acknowledge the research infrastructure support from the Department of Primary Industries (DPI), Queensland.

## Contributions

Conception and design were by Z.C, C.T.O, and E.M.R. Sample collection and experiments were by Z.C, C.T.O, M.R.S.F, K.M., M.H.D, A.C.B, and B.S.F. Data analysis and visualization were by Z.C, C.T.O, and E.M.R. Draft writing was by Z.C. Draft reviewing was by Z.C, C.T.O, M.R.S.F, K.M., M.H.D, A.C.B, B.S.F, and E.M.R.

